# Functional characterization of Lipid storage droplets 1 (LSD1) in growth and lipolysis of *Hermetia illucens*

**DOI:** 10.1101/2024.03.22.586280

**Authors:** Yuguo Jiang, Zongqing Kou, Bihui Chen, Luo Xingyu, Yongping Huang

**Affiliations:** CAS Key Laboratory of Insect Developmental and Evolutionary Biology, CAS Center for Excellence in Molecular Plant Sciences, Institute of Plant Physiology and Ecology, Chinese Academy of Sciences, Shanghai, China; University of Chinese Academy of Sciences, Beijing, China

**Keywords:** *Hermetia illucens*, perilipin, lipid homeostasis, CRISPR/Cas9, growth, fatty acid

## Abstract

As intracellular organelles in adipose tissue, lipids droplets manage the balance between triglyceride accumulation and energy consumption in animals. Perilipin family members, associated with surface of lipid droplets, participate the regulation of lipid metabolism. Lipid storage droplet-1 (LSD1)/Perilipin-1 acts as a gatekeeper for adipose lipid storage in animals. Despite extensive studies in fruit fly, the function of *LSD1* in insect larval stage remain indistinct. In this study, we characterized the function of *LSD1* in black soldier fly *Hermetia illucens*, a nova resource insect to recycle organic wastes. We found that *LSD1* was broadly present in dipteran species and evolved with divergence between mosquitos and flies. We further constructed *in vivo* mutagenesis mediated by CRISPR/Cas9 and found that mutations in *LSD1* increased the larval weight and did not bring any defects in development. Raw fat content was also not significantly influenced in late larval stage and new-emerged adults. Our results not only extend our knowledge of *LSD1* in insects, but also help for better understanding of the lipid homeostasis in BSF.

## Introduction

Lipid homeostasis, balance between lipogenesis, storage, and lipolysis, is fundamental for growth, development, and reproduction in living organisms. In insects, fat body, primarily composed of adipocytes, is the hub of lipid storage and metabolism (Li et al., 2019). As the core intracellular organelle in adipocytes, lipid droplets (LDs) play the critical role in lipid homeostasis. LDs consist of a monolayer of phospholipid with a triglyceride (TAG) core, which is proposed to be formed from endoplasmic reticulum (Walther et al., 2017). For lipogenesis, fatty acids, hydrolyzed form diet lipids or synthesized *de novo*, serve as acyl donors for the esterification. TAG, as the terminal product, will be transported into LDs for storage. To responded to the energy demand, TAG will be disintegrated back into glycerol and fatty acids by lipid lipases located on the surfaces of LDs and the fatty acids will go through continuous beta-oxidation to produce energy (Arrese & Soulages, 2010).

The accumulation of triglycerides in fat body tissues is a balanced results of the dynamics between neutral lipogenesis and lipolysis. For lipolysis, perilipins is the key factor to recruit lipases from cytoplasm to LDs (Brasaemle, 2007). In *Drosophila melanogaster*, two perilipins family members are encoded, *Lipid storge droplet-1(LSD1)/ Perilipin-1* and *LSD2/Perilipin-2* (Lu et al., 2001). *DmLSD1* is expressed during all ontogenesis stages (Beller et al., 2010) and the protein is exclusively located on the surface of LDs in larval fat body (Bi et al., 2012). Opposite the guardian function of DmLSD2 to obstruct lipolysis, DmLSD1 could stimulate lipolysis through adipokinetic hormone (AKH) signaling pathway or by recruitment of lipases to LDs surface(Kimmel & Sztalryd, 2016). In fruit fly, absence of *LSD1* suppressed both above mentioned mechanisms and increased adiposity in adults (Bi et al., 2012). In fruit fly, a few study mentioned the changes in glyceride accumulation in larval stage of *lsd1*. Although under fed condition, average area of single LD was significantly increased in *lsd1* larvae compared with wild type, the total glyceride was not changed in *lsd1* mutants except that under starvation condition (Bi et al., 2012). In other non-drosophila insects, the species-specific functions of perilipins in lipid accumulation or lipolysis have not been demonstrated.

The black soldier fly (BSF), *Hermetia illucens* (L.) (Diptera: Stratiomyidae) has attracted extensive attention as one of the most promising insects that are qualified in organic wastes recycling (Raksasat et al., 2020; Surendra et al., 2020; Kim et al., 2021). This species could efficiently assimilate various organic substrates into biomass composed of protein, fat, and chitin, which are valuable raw materials or ingredients for industries and animal husbandry (Huis et al., 2020; Hawkey et al., 2021; Niyonsaba et al., 2021). Great potentials are held in BSF for its nutritional value. For the inside grease, after extraction, separation, and refinement, could be added into feed (Fawole et al., 2021; Heuel et al., 2021) or cosmetic products (Almeida et al., 2020) as gradients, depending on the depth of refinement, or as raw materials for biodiesel (Zheng et al., 2012; Nguyen et al., 2017; Wang et al., 2017) and biolubricants (Xiong et al., 2020) after transesterification. BSF, as a nova resource insect, is worthy of conducting research on functions of perilipins in lipid homeostasis to maximize its advantages for application.

In our study, *HiLSD1* was cloned from genome of *H. illucens* and two homozygote mutants of *HiLSD1* were obtained based on CRISPR/Cas9 platform. The function role of *HiLSD1* in larval development was initially characterized by growth assay. Besides the influences on lipid droplets forming, we also investigated whether and how *Hilsd1* change the fat accumulation and fatty acids profiles. The results will not only extend our knowledge of *LSD1*, but also help for better understanding of lipid metabolism in BSF.

## Materials and Methods

### 1. Rearing of BSF strains

The laboratory-maintained wild-type line of BSF, which was originally collected from Dr. Ziniu Yu’s lab, underwent inbred crossing for generations in our lab (Zhan et al., 2019). The newly hatched larvae, wild type, or mutants were reared in moistened wheat bran (1:2 ratio of dry matter/ddH_2_O) under an 8:16 L:D photoperiod at 28℃ with relative humidity of 50%-60%.

### 2. Clone of *LSD1* in BSF

The amino acid sequence of LSD1 in *D. melanogaster* (NP_001262883.1) was used as a query to search the ortholog in the genome of the BSF (GCF_905115235). Based on manual annotations, the architecture of ORF were predicted. Polymerase chain reaction (PCR)-amplification was applied to obtain coding sequence from complementary DNA (cDNA) with the following pair of primers: forward, 5’-ATGGTTAAGCAACAGATAAAACG-3’ and reverse, 5’-TCAATAAACTCCATTTAAATTGTTC-3’. For cDNA synthesis, total RNA was isolated from 3^rd^ larval instar of wild-type strain using TRIeasyTM Total RNA Extraction Reagent (Yeasen, Shanghai, China). One µg of total RNA was processed using PrimeScript RT reagent Kit with gDNA Eraser (Takara, Beijing, China). The PCR procedure was set as follows: 98℃ for 2 min, followed by 30 cycles at 98℃ for 30 s, 55℃ for 30 s, 68℃ for 1 min, and an elongation phase at 68℃ for 10 min. Amplified products were sequenced and verified after cloning into a pJET1.2-T vector through CloneJET PCR Cloning Kit (Thermo Scientific, Waltham, USA).

### 3. Comparative and phylogenetic analysis of LSD1

The protein sequence of HiLSD1 was used as a query to search dipteran orthologs against Genbank nonredundant protein database (nr) using BLASTP. The sequences from 8 representative dipteran species were used for further analysis (*Aedes aegypti*, XP_021693334.1; *Anopheles gambiae*, XP_312022.5; *Culex quinquefasciatus*, XP_001845547.2; *Bactrocera dorsalis*, XP_011212558.1; *Ceratitis capitata*, XP_004523385.1; *Drosophila melanogaster*, NP_001262883.1; *Lucilia cuprina*, XP_023297473.1; *Musca domestica*, XP_005189534.1;) with a lepidopteran species as reference (*Manduca sexta*, ACF24761.1). Protein sequences of HiLSD1 and orthologous sequences were subjected to multiple alignment and phylogenetic analysis using MEGA X (Li et al., 2022). The phylogenetic tree was inferred using the maximum likelihood method with a bootstrap of 1000 replications. The tree was rooted using the ortholog of *Manduca sexta* as the outgroup.

### 4. CRISPR/Cas9-mediated construction of *lsd1* homozygotes

For CRISPR/Cas9-mediated mutagenesis of *LSD1*, with the PAM sequences in consideration, newly designed sgRNAs should follow the NNN19GG rule (Zhan et al., 2019). Based on sequence identification of *HiLSD1*, four 23-bp sgRNA targeting sites named S1-4 (Table S1) were designed by Geneious Prime 11.0.3. sgRNA templates were transcribed using a T7 promoter and synthesized *in vitro* using the MAXIscript T7 kit (Ambion, Austin, USA) according to the manufacture’s instruction. The Cas9 protein was purchased form Thermo Scientific.

Germline transformation was performed as described previously (Zhan et al., 2019). Briefly, Cas9 protein (200 ng/µL) with two sgRNAs (100 ng/µL) were co-injected into embryos. After hatching, first instar larvae were selected to detect genomic mutations with primers listed in Table S2. The amplified fragments were cloned into pJET1.2 for sequencing. Frame-shift mutants were reared in moistened wheat bran for inbreed crossing to obtain homozygotes.

### 5. Growth assay of *lsd1*

To verify the effect of *lsd1* on larval growth, 4^th^ day larvae, wild type and *lsd1*, were reared in a chamber with temperature, relative humidity and fluorescent photoperiod set to 28 ℃, 60% RH and 8:16 (L:D). Sixty larvae of each genotype were placed into individual plastic basin (500 mL) with three replicates. One hundred gram of moistened wheat bran (1:2 ratio of dry matter/ddH_2_O) were refreshed every 24 hours until over 75% of larvae reached prepupal stage. Before routine reloading, weight of all larvae in each basin was recorded, as well as the number of survived and development stage at that very moment. The larval stage was determined by mouth morphology characteristics (2015-Kim). The larvae collected at 7 days, 12 days and 22 days post hatching were defined as L4, L5 and prepupae, respectively. All collected larvae for further chemical analysis were stored in–80℃ after inactivation by liquid nitrogen.

### 6. Feed experiment

The feeding trials, starting with 7^th^ day larvae (around the middle of L4), were under the same rearing condition as mentioned above. Sixty larvae were fed twice, being on days 1 and 4, with 120 g moistened wheat bran. The initial weight was monitored by weighing 20 larvae for each sample. At the end of trials on day 12, larvae were separated from the remaining residues and weighed. The dry matter content of residues and larvae was determined after dried in oven.

Bioconversion efficiency (BE), feed conversion ratio (FCR), waste reduction (WR) and efficiency of conversion of digested feed (ECD) were calculated as previously reported (Broeckx et al., 2021; Ribeiro et al., 2022):

BE = (DW_larvae, end_ – DW_larvae, start_) / DW_feed_ x 100%

FCR = DW_feed_ / (FW_larvae, end_ – FW_larvae, start_)

WR = (DW_feed_ – DW_residue_) / DW_feed_ x 100%

ECD = (DW_larvae, end_ – DW_larvae, start_) / (DW_feed_ – DW_residue_) x 100%

Where DW is the dry weight and FW is the fresh weight.

### 7. Lipid droplet staining and microscope

Fat body tissue from 11-day-old larvae (middle 5^th^ instar stage) was prepared as follows for *ex vivo* confocal laser scanning microscopy. The body of 5^th^-instar larva was opened, and the fat body tissues attached to the cuticle mechanically released into mounting medium (50% glycerol/PBS, Nile red [Sigma, stock 10% in DMSO] 1:5000, Hoechst 33342 [Beyotime, stock in 10 mg/mL] 1:1000). The tissue was analyzed within 2h after mounting using a Leica Stellaris 5 with 405 nm excitation/420-471 nm emission wavelength range for Hoechst and 561 nm excitation/570-630 nm emission wavelength range for Nile red.

### 8. Quantification of raw fat content

All collected larvae or adults for chemical analysis were dried in an oven at 60℃ for 72h and reweighed to determine water content. The total fat content of BSF was determined using the standard protocol (GB/T 5009.6-2016). Briefly, 1 g of the dried sample was ground and subjected to Soxhlet extraction with n-hexane(AR, Sinopharm) at 75℃ overnight. After vacuum evaporation, the extracts were weighed to determine the raw fat content.

### 9. Analysis of fatty acid profiles

The fatty acid profiles of extracted raw fat were measured using the standard method (GB/T 5009.168-2016). Briefly, 50 mg sample was put into 5 mL glass vial with 1 mL of NaOH-MeOH solution (2%, w/v) for saponification. To obtain the methyl ester solution, 1mL BF_3_-MeOH (14%, sigma) was added into the vial for 15 min, followed with 2 mL n-hexane to extract the products. Finally, the fatty acid methyl esters (FAMEs) were determined by gas chromatography.

An GC2010 pro (Shimazu, Suzhou, China) was equipped with a TG-WaxMS capillary column (60 m × 0.25 mm × 0.25 µm, Thermo scientific) and a flame ionization detector (FID). The GC program was as follows: initial temperature of 120 °C; ramp to 240℃ at rate of 5℃/min, maintain 6min. The temperature of inlet and detection were 220℃ and 240℃, respectively. The sample volume of per injection was 2 µL in split ratio of 40. The flow rate of carrier gas (nitrogen) was kept at 1.5 mL/min. Qualitative analysis was conducted by authentic compounds purchased from Sigma. Fatty acid relative content was quantified by percentage of weight of each fatty acid in raw fat, which was calculated by external standard method.

### 10. Investigation of expression of relevant genes

Relative expression of several genes in insulin signaling pathway and lipid homeostasis network from 11-day-old larvae (middle 5^th^ instar larval stage) was determined in *lsd1* mutant and WT. Total RNA was extracted from fat body of larvae and subsequently reverse-transcribed into cDNA as mentioned above. The transcript levels were assayed using SYBR Green Master Mix (TOYOBO) normalized against *HiActin* (Gao et al., 2019). The primers are listed in Table S3.

### 11. Statistics

A two tail Student’s t-test was used to analyze the differences between results unless otherwise noted. Bar charts were drawn and analyzed with GraphPad PRISM 8.0 software (San Diego, California, USA). Three independent replicates were used for each test. Error bars represent the standard deviation (S.D.).

## Results

### 1. Characterization of *HiLSD1* in the genome

To identify sequence of *HiLSD1*, the protein sequence of *LSD1* in *D. melanogaster* (NP_001262883.1) was used to characterize the ortholog (*HiLSD1*) in genome of *Hermetia illucens* (GCF_905115235) (Generalovic et al., 2021). *HiLSD1* is located on chromosome 1 and spans 14433-bp genomic sequence, which consists of 7 exons and encodes a 398-amino acid protein (Fig. 1A). Using *HiLSD1* as a query for BLAST, we found that most dipteran insect, especially representative flies, and mosquitos, encoded *LSD1* in their genome, but nevertheless shares a relatively low amino acid identity (from 54.50% in *Bactrocera dorsalis* to 57.18% in *Drosophila melanogaster*). In addition, the phylogenetic analysis of dipteran *LSD1* orthologs showed that the topology of the gene tree greatly agreed with that of the species tree by previous study (2019-CR)(Fig. 1B), which implied that the evolution of *LSD1* may correlate with the speciation across dipterans. To conclude, above results reveal that LSD1 is broadly present in dipterans and under conserve evolution across species.

**Fig. 1.**
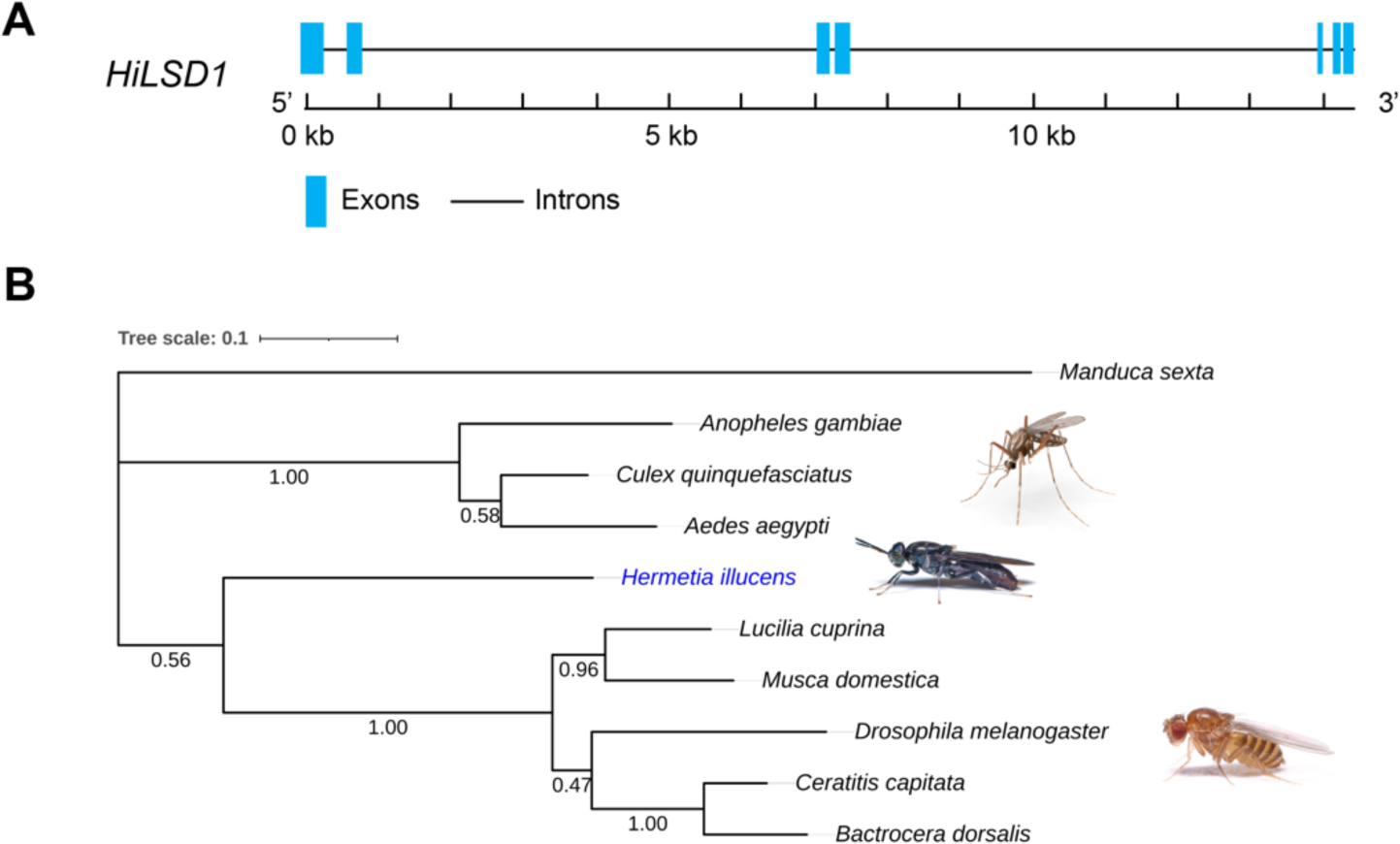
The genome structure of *HiLSD1* and evolutionary relationship among representative dipteran insects. (A) Schematic of *HiLSD1* gene. The blue boxes represent exons, and the solid lines represent introns. (B) Phylogenetic analysis of HiLSD1. The analysis involved 10 amino acid sequences, using LSD1 from *M. sexta* as outgroup. *Hermetia illucens* is highlighted in blue.

### 2. Mutagenesis of *Hillsd1* mediated by CRISPR/Cas9

To further investigate the function of HiLSD1 in BSF, an *in vivo* knock-out was conducted mediated by CRISPR/Cas9 system (Fig. 2A). Two pairs of sgRNAs, Target site 1/ Target site 2 (TS1/TS2) and Target site 3/ Target site 4 (TS3/TS4) were designed in preceding exons following the NNN19GG rule (Fig. 2B). The transformation was performed on fresh laid eggs with microinjection equipped with 200 ng/µL Cas9 protein and 100 ng/µL for each paired sgRNAs. Newborns of G1 were selected to detect the mutations. Positive mutants were reared generations for inbreed crossing to obtain homozygotes (Fig. 2A). Both homozygotes shown in Fig. 2B were obtained in G3 offspring after mutation detection for over 10 egg clutches in each generations. Both mutants showed disruption in amino acid sequences of HiLSD1 and caused premature translational termination, indicative of at least partial function loss. It should be noted that design of pair sgRNAs located in exons was for large fragment deletion, which did not happen(Fig. 2B). Adjustment of Cas9 and sgRNAs concentration, along with selection of sgRNAs for higher theoretical efficiency, would increase the chance for large fragment deletions.

**Fig. 2.**
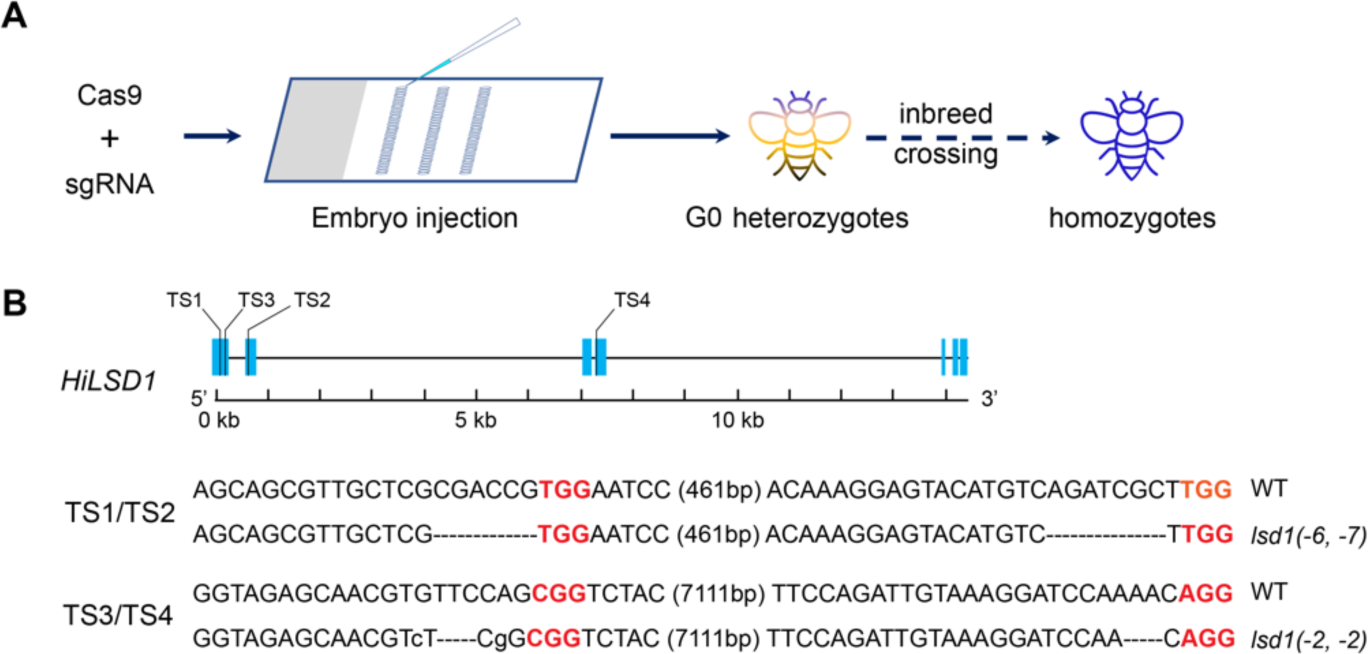
CRISPR/Cas9 mediated mutagenesis of *HiLSD1* for loss-of-function homozygotes. (A) Graphic workflow of *Hilsd1* construction. Colorful fly represents heterozygotes, and blue one represents homozygotes. (B) Schematic mutagenesis target sites (TS) of *HiLSD1* and genotype of obtained homozygotes. PAM sequences are shown in red and TS sequences are in bold black. Dashed lines indicate deleted base pairs and lowercase letters mean mutated base pairs. Number of deleted base pairs is shown in the brackets.

### 3. Improvement in larval growth by *Hilsd1*

To investigate the influence in growth by *Hilsd1*, a growth assay was conducted among two mutants and wild type (WT). For incubation, similar size and average weight was considered to control the initial larval conditions. Daily refreshed moistened wheat bran as food is guaranteed to be sufficient. For larval weight during the development, both *lsd1* mutants achieved significant heavier body than WT (Fig. 3A). For WT, maximum average weight, 0.2210 g, appeared at 269h (around 11 days after hatching), which was late stage of 5^th^ instar larvae. Then the average weight decreased, and the larvae began to pre-pupate. Meanwhile, both *lsd1* mutants were still in weight growth, 0.2416 g of *lsd1*(–6,-7) and 0.2320 g of *lsd1*(–2,–2), which were 9.30% and 4.98% heavier than WT, respectively. Maximum average weight of *lsd1* reached at 295h, the time that prepupation had begun in all groups. The average weight of *lsd1*(–6,-7) was 0.2504 g and that of *lsd1*(–2,-2) was 0.2400 g, which were 15.67% and 10.85% heavier than WT with little decreased weight (0.2165 g) at that timepoint, respectively. After that, weight of larvae in all groups decreased caused by prepupation and pupation. Both mutants maintained a significant advantage in weight until 18 days after hatching (around 440h). At the end of pupae stage, when the first eclosion appeared, rest of all pupae were weighed and pictured. For *lsd1*(–6,-7), mean of pupae weight was significantly little heavier than WT (7.24%, WT: 0.1380 g; *lsd1*(–6,-7): 0.1480 g) (Fig. 3B).

**Fig. 3.**
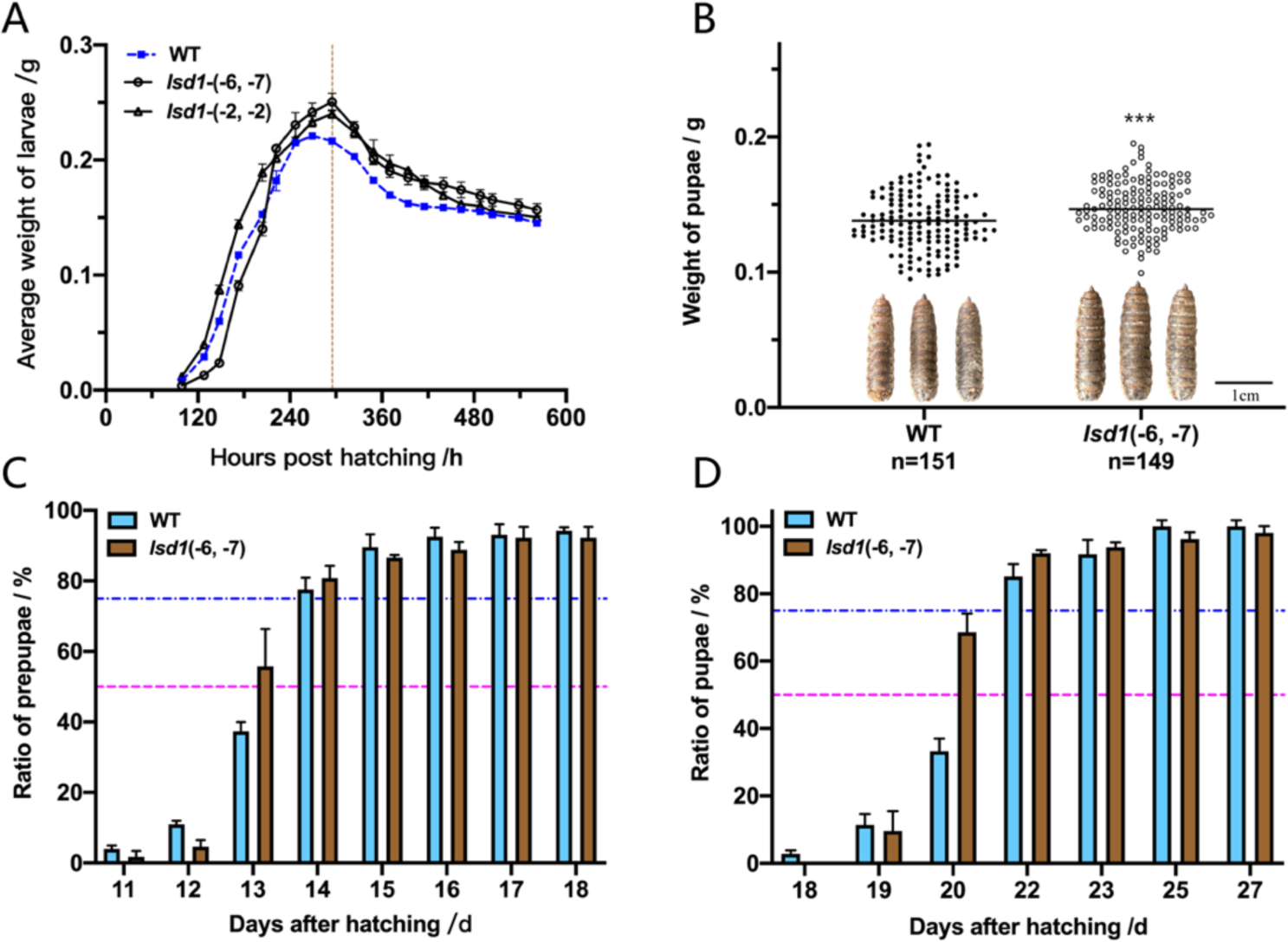
Phenotype of *Hilsd1* in larval growth and development. (A) Changes in the larval weight during the growth until late pupae stage. Blue dashed line indicates wild type (WT) while hollow circle and triangle represents *lsd1*(–6,–7) and *lsd1*(–2,–2), respectively. Vertical dashed line at 295h points the maximum weight of mutants. (B) Pupae weight of WT and *lsd1.* Solid lines across the dots indicated the mean of the weight. Differences of pupae weight between WT and *Hilsd1* were tested by Welch’s test. (C) Prepupation and pupation (D) rate of WT and *lsd1.* Magenta dashed line points at 50% and the blue one is at 75%.

For development, the mutant would little early reach 50% prepupation (Fig. 3C) and 50% pupation (Fig. 3D), especially in a situation that first prepupae and pupae always appeared in WT group. As illustrated, pupae of *lsd1*(–6,-7) were little larger than that of WT. Taken together, disruption of *HiLSD1* did not bring any apparent defects in larval growth and development, even improve the growth in larval stage instead.

### 4. The better feeding performance of *Hilsd1*

To further elucidate the reason of improved growth in *Hilsd1*, a feeding trial was performed during L4-L5 (7 day – 12 day) which cover the steep part of growth curve (Fig. 3A). The waste reduction (WR) and ECD were displayed in Table 1. These two parameters are good indicators of feed conversion efficiency. Though no significance in WR between WT and mutant, over 10% increase in ECD of *lsd1*, compared with WT, suggested a better assimilation of moistened wheat bran in mutant. The lower value of FCR and higher value of BE in *lsd1* also indicated mutants would show better feeding performance and grow bigger or faster than WT. As mentioned above, *Hilsd1* grew a little heavier (over 10%) than WT around 12^th^ day after hatching (Fig. 3A), which would be contributed by the higher assimilation efficiency of *Hilsd1*.

**Table. 1.** Comparison of Bioconversion efficiency, feed conversion ratio, waste reduction and ECD between *lsd1 (–6,-7)* and WT. Values are reported as mean ± standard deviation (replicates indicated in the table as *n*). ***, p < 0.001; ns, no significance according to Student’s *t* test.

|  | *n* | Bioconversion Efficiency (%) | Feed Conversion Ratio (FCR) | Waste reduction (%) | Efficiency of Conversion of Digested feed (ECD) |
| --- | --- | --- | --- | --- | --- |
| WT | 6 | 3.47 ± 0.10 | 11.08 ± 0.70 | 44.20 ± 0.34 | 7.86 ± 0.26 |
| *lsd1*(-6, -7) | 6 | 3.88 ± 0.07*** | 8.44 ± 0.25*** | 43.49 ± 1.52<sup>ns</sup> | 8.92 ± 0.37*** |

### 5. The fusion lipid droplets in larvae of *HiLSD1* mutation

As mentioned in previous study (Kimmel & Sztalryd, 2016), LSD1 in *Drosophila melanogaster* located on surface of lipids droplets and was involved in lipid droplets (LDs) fission or fusion to regulate neutral lipids metabolism. The fat body of 5^th^-instar larvae, WT and *lsd1*(–6,–7), were dissected from the tissues attached to the cuticle and released into a mounting medium containing Hoechst for nucleus staining and Nile red for neutral lipids staining. As illustrated, consistent with study in *D*. *melanogaster* (Bi et al., 2012), *Hilsd1* mutant larval fat bodied had apparently larger lipid droplets than WT (Fig. 4). Given that a complete N-terminal of LSD1 (112 amino acid) is kept in *lsd1*(–6,–7) due to the target site position, which is predicted to be the N-terminal of a perilipin domain by Conserved Domains Search (https://www.ncbi.nlm.nih.gov/Structure/cdd/wrpsb.cgi), there were incomplete LD fusion in *lsd1*(–6,–7) as illustrated.

**Fig. 4.**
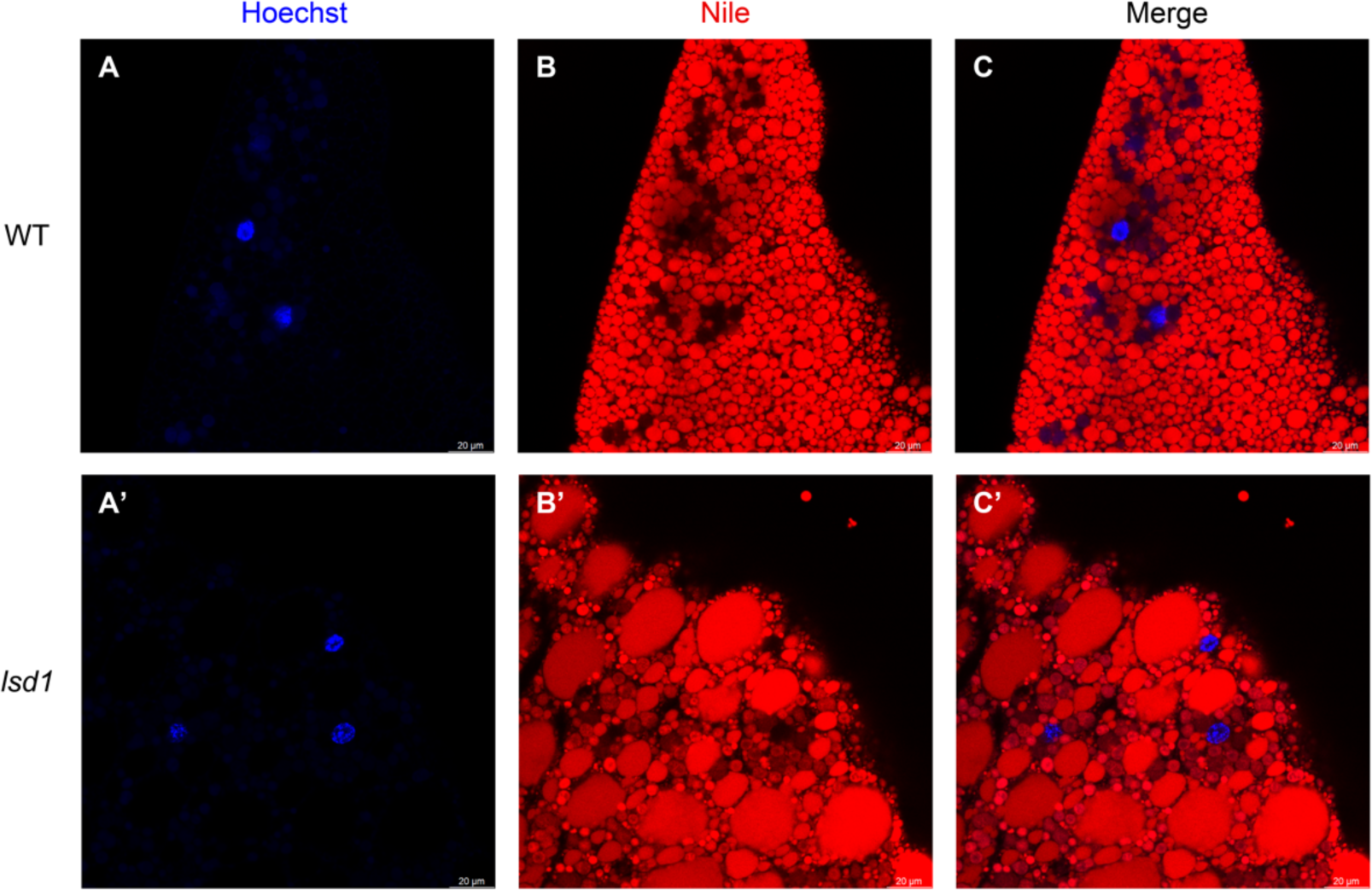
Cellular effect of *Hilsd1* on lipid droplets in 5^th^-instar larvae’s fat body. Nucleus (blue) and neutral lipids (red) were simultaneously stained with Hoechst and Nile red, respectively.

### 6. The impacts of *Hillsd1* in raw fat content and fatty acids profiles

As LDs fusion may influence the neutral lipids degradation, the raw fat content was quantified by Soxhlet extraction with non-polar solvent n-hexane. Although *lsd1* mutation promoted LDs fusion, raw fat content per larvae and percentage of dry materials did not show any significant difference between WT and *lsd1* in 4^th^-instar larvae, 5^th^-instar larvae and prepupae stage (Fig. 5AB). However, according to the content per larvae, *Hilsd1* appeared to show a tendency for more fat accumulation (Fig. 5A). For fatty acid profiles (Fig. 5C), no significant change was brought by *lsd1* mutation.

**Fig. 5.**
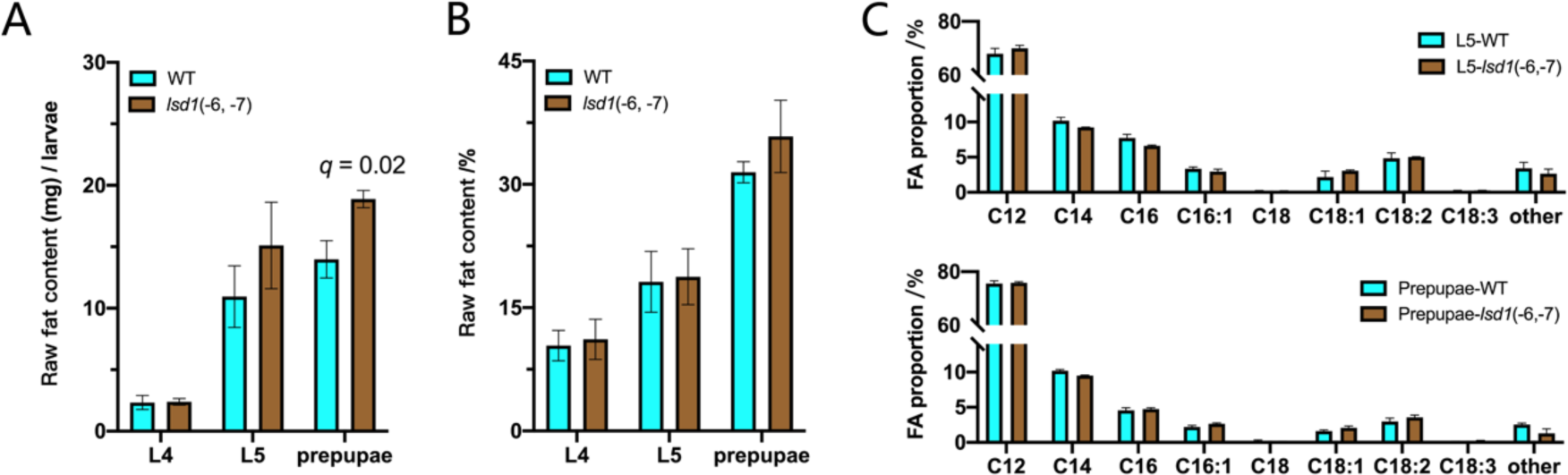
The impacts of *Hilsd1* on raw fat content and fatty acid profiles. (A) Raw fat content per larvae or percentage of dray weight (B) and fatty acid (FA) profiles (C) of WT and *Hilsd1* in different stages: 4^th^ & 5^th^-instar larvae and prepupae.

### 7. Investigation of lipogenesis relevant genes in *Hillsd1*

To further investigate the cause of the phenotype by impaired HiLSD1, several genes in insulin signaling pathway, lipolysis pathway and fatty acid metabolism pathway were selected to detect the changes in expression level in 5^th^-instar larvae’s fat body (Fig. 6). For insulin signaling pathway, the expression of all selected genes, except *InR*, significantly upregulated in *Hilsd1* in comparison to WT, which partially illustrate the size change around 5^th^-instar stage (Fig. 6A). For TAG degradation, we expected that *Hilsd1* mutation would upregulate *HiLSD2* like drosophila (Bi et al., 2012) and activate *HSL* and *Bmm* for more TAG triglyceride. Unexpected, all the three genes had no significant change which indicate a different regulation mechanism from fruit fly (Fig. 6B). For fatty acid metabolism, *FAS* for *de novo* lipogenesis was regulated (Fig. 6C), while fatty acid degradation pathway were downregulated by less acyl-CoA provision from upstream, which had been prevented by *lsd1* mutant. In fatty acid profiles of WT and *Hilsd1*, from 5^th^-instar larvae to prepupae stage, changes in relative percentage in lauric acid (C12) and myristic acid (C14) could not show the effect of the upregulated *de novo* lipogenesis for existing high content. The smaller decreased margin of palmitic and palmitoleic acid content in *Hilsd1* illustrated the downregulated fatty acid degradation pathway.

**Fig. 6.**
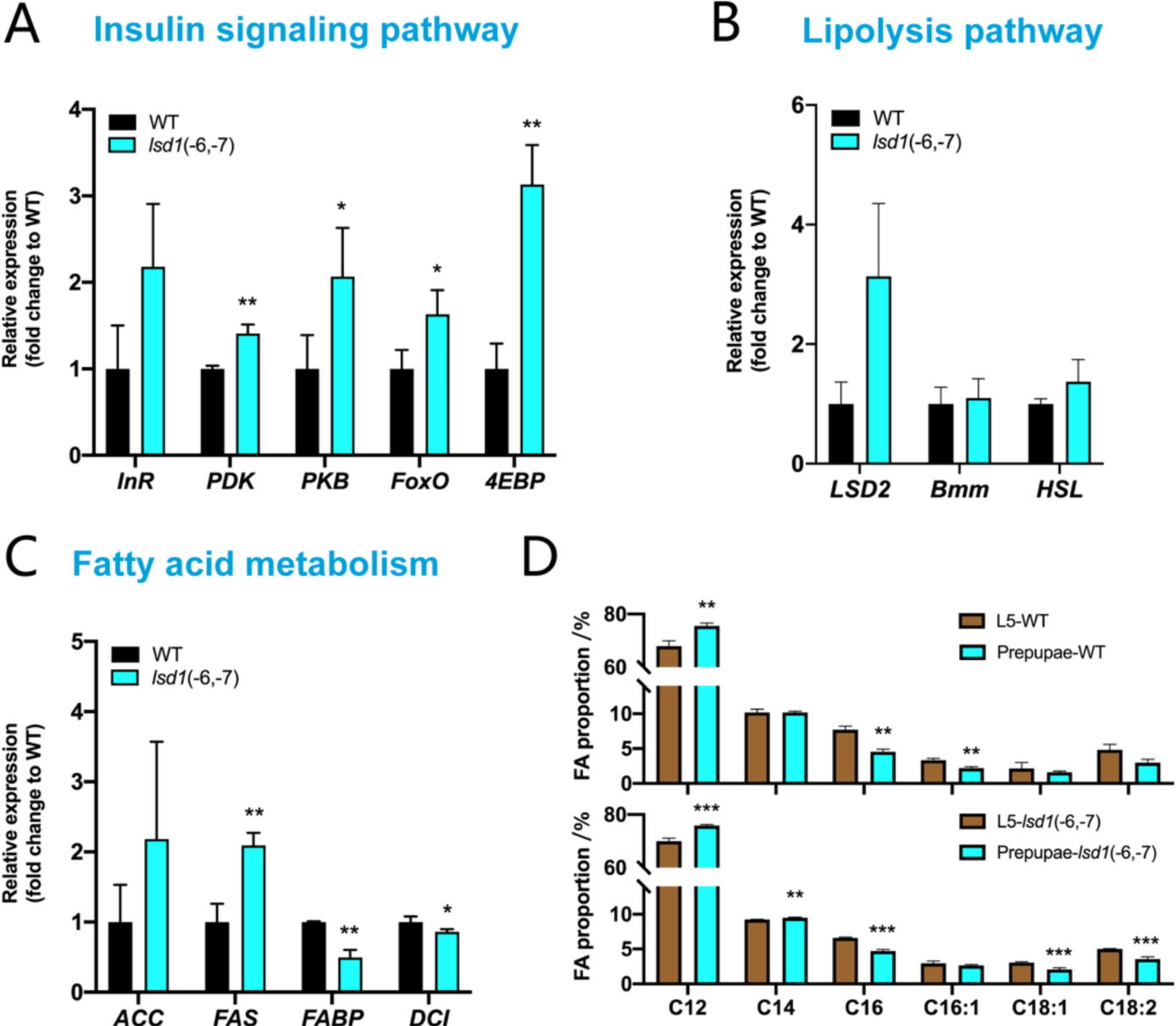
Changes in mRNA levels of relevant pathway and fatty acid profiles in growth. The relative mRNA levels of relevant genes in fat body of the 5^th^-instar larvae were determined using *Actin* as a control. (A) *InR*, *PDK*, *PKB, FoxO* and *4EBP* were chosen to detect changed in insulin signaling pathways; (B) *LSD2*, *Bmm* and *HSL* were to detect the impact in lipolysis pathway; (C) *ACC, FAS*, *FABP* and *DCI* were for fatty acid metabolism pathway. (D) Changes in fatty acid profiles from L5 to prepupae in WT and *lsd1*. The data are shown as the mean ± S.D. (n=3). *, p < 0.05; **, p < 0.01; ***, p < 0.001 according to Student’s *t* test.

## Discussion

LSD1 is a member of perilipin family, which is associated with lipid accumulation and energy consumption in lipid droplets (Kimmel & Sztalryd, 2016).The function of LSD1 has been extensively studied in murine tissues and adult flies, but the functions in insect larval growth and development have been less addressed. In larvae stage of drosophila, *lsd1* brought increase in relative glyceride levels only in starved condition, while no significant changes were appeared in feeding condition (Bi et al., 2012). In addition, the effect of growth and development may be too little to be noticed in drosophila larvae. For BSF, given the main commercialized products were from larval stage, including prepupae, the mutation effect on the growth, development and glyceride levels would be more worthy of attention. In this study, we first characterized the ortholog of *LSD1* in *H. illucens* and uncovered a widespread distribution in representative dipteran species (Fig. 1). Compared with LSD1 from *M. sexta*, orthologs of LSD1 in dipteran species were relatively conserved. Compared with phylogeny relationship of species (Zhan et al., 2019), evolution of LSD1 may correlate with the speciation across dipterans.

To investigate the function of *HiLSD1*, CRIPSR/Cas9 knock-out system was employed to generate mutations *in vivo*. After inbreed crossing for several generations, two homozygote mutants were obtained (Fig. 2). Despite of predicted premature translational termination, *Hilsd1* showed no significant defects in growth and development (Fig. 3). The mutant adults behaved as the same vigor as WT, which was observed by naked eyes in short distance. Instead, mutants, especially *lsd1*(–6,-7), grew into heavier larvae than WT in late 5^th^-intar stage, beyond 10% (Fig. 3A). Even after pupation with weight decreasing, *lsd1* still kept the advantage in weight, which was an unexpected results due to the phenotype of lean body but with equal weight in mice (Tansey et al., 2001). By feeding assays, *Hilsd1* showed significantly increased assimilation than WT (Table 1), which correspond with the heavier body of mutants.

As the same to drosophila, *Hilsd1* also caused lipid droplets fusion (Fig. 4) which indicated inhibition for lipolysis. Although no significance increase was brought in raw fat content by *lsd1* (Fig. 5A), but the tendency suggested by the changes in content per larvae implied that the relevant genes in lipid homeostasis network had been stroke by mutations in *lsd1*. Further investigation in gene expression pattern showed that insulin signaling pathway had been upregulated in *Hilsd1* mutants, which would partially explain the bigger size and obviously heavier larval weight. Lipolysis pathway was not impaired significantly by mutations, while fatty acid metabolism was affected to weaken the downstream fatty acid degradation (Fig. 5C), which contributed to the abnormal raw fat accumulation in *Hilsd1* prepupae stage.

In summary, this study initially investigated the *in vivo* function of *LSD1* in *H. illucens* and focused on its role in larval stage. The findings not only revealed the capacity of *HiLSD1* in lipid homeostasis and insect development, but also shed a light on the lipogenesis for lauric acid through inhibition of fatty acid oxidation by *lsd1*, which would lead to the elucidation of the uncommon fatty acid profiles in BSF.

## Acknowledgement

The authors would like to thank the financial support for this work by the Shanghai Pudong New Area Science and Technology Development Fund (PKJ2019-C01).

