## supplemental materials for "Functional characterization of Lipid storage droplets 1 (LSD1) in growth and lipolysis of *Hermetia illucens*"

This profile includes:

Table S1-3

**Table S1: sgRNA target sites (TSs) for mutation construction**

| TS names | TS sequence, 5’->3’ |
|---|---|
| TS1 | AGCAGCGTTGCTCGCGACCGTGG |
| TS2 | GGAGTACATGTCAGATCGCTTGG |
| TS3 | GGTAGAGCAACGTGTTCCAGCGG |
| TS4 | GATTGTAAAGGATCCAAAACAGG |

**Table S2: Primers for mutation detection**

| Primer name | Primer sequence, 5’->3’ |
|---|---|
| TS1-det-Fw | GCAACATCCCTATCATTGAGACA |
| TS1-det-Rv | AGGGAGATATAATCCCAATGCCA |
| TS2-det-Fw | AGGTTCACTAAGTTCTTCACGTC |
| TS2-det-Rv | GATCTGCGTTTCCTTCGACT |
| TS3-det-Fw | GCAACATCCCTATCATTGAGACA |
| TS3-det-Rv | AGGGAGATATAATCCCAATGCCA |
| TS4-det-Fw | CGAACTATCTTGGAAGCACG |
| TS4-det-Rv | TCTTGATTTACCTGGGAATGC |

**Table S3: Primers for qPCR**

| Primer name | Primer sequence, 5’->3’ |
|---|---|
| FAS-Fw | AAACCTCCAATCTTCTTCGTGC |
| FAS-Rv | CAAGTGGTGCCTCTTCTGTGC |
| ACC-Fw | AACAGTGGTGCTCGTATC |
| ACC-Rv | TGGCTCGTCTTCATCTTC |
| FABP-Fw | GAATACATGAAGGCGCTCGG |
| FABP-Rv | CGGCTTTGAATTTGATGGCG |
| DCI-Fw | GGTTGTGGTGCTAACAGGTG |

| | |
|---|---|
| DCI-Rv | GGTCCATTGATGACTGCCAC |
| AKT-Fw | GAATATCTAGCGCCCGAGGT |
| AKT-Rv | CATCGTGGTCGCTGTTGTAG |
| PDK-Fw | GTACATCGCGACCTGAAACC |
| PDK-Rv | TTCGAATAAGAAGCTGCCGC |
| 4EBP-Fw | ATGCTACTGCCATCCCAACT |
| 4EBP-Rv | TTCTTCGGTGGGGTTTGACT |
| Bmm-Fw | TCCTCGGCATCTACCATGTC |
| Bmm-Rv | GGTCCAAGTGAATGACGACG |
| HSL-Fw | TTGGTCCGATGGCAGTTACT |
| HSL-Rv | TGTGACTTTGAACTCTGCGC |
| LSD2-Fw | TTAGCGGAGTGGACCAAACT |
| LSD2-Rv | ACTTTGTTGGAAAGACGCCC |
| Actin-Fw | CGTAGGAGACGAAGCACAAA |
| Actin-Rv | GGTGCCAGATCTTCTCCATATC |
